# Transmission dynamics of Zika virus in island populations: a modelling analysis of the 2013-14 French Polynesia outbreak

**DOI:** 10.1101/038588

**Authors:** Adam J. Kucharski, Sebastian Funk, Rosalind M. Eggo, Henri-Pierre Mallet, W. John Edmunds, Eric J. Nilles

## Abstract

Between October 2013 and April 2014, more than 30,000 cases of Zika virus (ZIKV) disease were estimated to have attended healthcare facilities in French Polynesia. ZIKV has also been reported in Africa and Asia, and in 2015 the virus spread to South America and the Caribbean. Infection with ZIKV has been associated with neurological complications including Guillain-Barré Syndrome (GBS) and microcephaly, which led the World Health Organization to declare a Public Health Emergency of International Concern in February 2015. To better understand the transmission dynamics of ZIKV, we used a mathematical model to examine the 2013–14 outbreak on the six major archipelagos of French Polynesia. Our median estimates for the basic reproduction number ranged from 2.6–4.8, with an estimated 11.5% (95% CI: 7.32–17.9%) of total infections reported. As a result, we estimated that 94% (95% CI: 91–97%) of the total population of the six archipelagos were infected during the outbreak. Based on the demography of French Polynesia, our results imply that if ZIKV infection provides complete protection against future infection, it would take 12–20 years before there are a sufficient number of susceptible individuals for ZIKV to reemerge, which is on the same timescale as the circulation of dengue virus serotypes in the region. Our analysis suggests that ZIKV may exhibit similar dynamics to dengue virus in island populations, with transmission characterized by large, sporadic outbreaks with a high proportion of asymptomatic or unreported cases.

**Author Summary:** Since the first reported major outbreak of Zika virus disease in Micronesia in 2007, the virus has caused outbreaks throughout the Pacific and South America. Transmitted by the *Aedes* species of mosquitoes, the virus has been linked to possible neurological complications including Guillain-Barre Syndrome and microcephaly. To improve our understanding of the transmission dynamics of Zika virus in island populations, we analysed the 2013–14 outbreak on the six major archipelagos of French Polynesia. We found evidence that Zika virus infected the majority of population, but only around 12% of total infections on the archipelagos were reported as cases. If infection with Zika virus generates lifelong immunity, we estimate that it would take at least 15–20 years before there are enough susceptible people for the virus to reemerge. Our results suggest that Zika virus could exhibit similar dynamics to dengue virus in the Pacific, producing large but sporadic outbreaks in small island populations.

## Introduction

Originally identified in Africa [1], the first large reported outbreak of Zika virus (ZIKV) disease occurred in Yap, Micronesia during April–July 2007 [2], followed by an outbreak in French Polynesia between October 2013 and April 2014 [3], and cases in other Pacific countries [4, 5]. During 2015, local transmission was also reported in South American countries, including Brazil [6, 7] and Colombia [8].

Transmission of ZIKV is predominantly vector-borne, but can also occur via sexual contact and blood transfusions [9]. The virus is spread by the *Aedes* genus of mosquito [10], which is also the vector for dengue virus (DENV). ZIKV is therefore likely to be capable of sustained transmission in other tropical areas [11]. As well as causing symptoms such as fever and rash, ZIKV infection has also been linked to increased incidence of neurological sequelae, including Guillain-Barré SYNDROME (GBS) [12, 13] and microcephaly in infants born to mothers who were infected with ZIKV during pregnancy [14]. On 1st February 2015, the World Health Organization declared a Public Health Emergency of International Concern in response to the clusters of microcephaly and other neurological disorders reported in Brazil, possibly linked to the recent rise in ZIKV incidence. The same phenomena were observed in French Polynesia, with 42 GBS cases reported during the outbreak [13, 15]. In addition to the GBS cluster, there were 18 fetal or newborn cases with unusual and severe neurological features reported between March 2014 and May 2015 in French Polynesia [16], including 8 cases with microcephaly [17].

Given the potential for ZIKV to spread globally, it is crucial to characterize the transmission dynamics of the infection. This includes estimates of key epidemiological parameters, such as the basic reproduction number, *R*_0_, defined as the average number of secondary cases generated by a typical infectious individual in a fully susceptible population, and how many individuals (including both symptomatic and asymptomatic) are typically infected during an outbreak. Such estimates could help assist with outbreak planning, assessment of potential countermeasures, and the design of studies to investigate putative associations between ZIKV infection and other conditions.

Islands can be useful case studies for outbreak analysis. Small, centralized populations are less likely to sustain endemic transmission than a large, heterogeneous population [18], which means outbreaks are typically self-limiting after introduction from external sources [19]. Further, if individuals are immunologically naive to a particular pathogen, it is not necessary to consider the potential effect of pre-existing immunity on transmission dynamics [20]. Using a mathematical model of vector-borne infection, we examined the transmission dynamics of ZIKV on six archipelagos in French Polynesia during the 2013–14 outbreak. We inferred the basic reproduction number and the overall size of the outbreak, and hence how many individuals would still be susceptible to infection in coming years.

## Methods

### Data

We used weekly reported numbers of suspected ZIKV infections from the six main regions of French Polynesia between 11th October 2013 and 28th March 2014 (Table 1), as detailed in the Centre d’hygiène et de salubrité publique situation reports [21, 22]. Confirmed and suspected cases were reported from sentinel surveillance sites across the country; the number of such sentinel sites varied in number from 27–55 during the outbreak (raw data are provided in S1 Dataset). Clinical cases were defined as suspected cases if they presented to health practitioners with rash and/or mild fever and at least two of the following signs: conjunctivitis, arthralgia, or oedema. Confirmed cases were defined as a suspected case if they tested positive by RT-PCR on blood or saliva. In total, 8,744 suspected cases were reported from the sentinel sites. As there were 162 healthcare sites across all six regions, it has been estimated that around 30,000 suspected cases attended health facilities in total [21]. For each region, we calculated the proportion of total sites that acted as sentinels, to allow us to adjust for variation in reporting over time in the analysis. Population size data were taken from the 2012 French Polynesia Census [23]. In our analysis, the first week with at least one reported case was used as the first observation date.

**Table 1.** Geographical breakdown of the 2013–14 French Polynesia ZIKV outbreak.

| Regions | Population | Suspected cases | PCR confirmed cases |
| --- | --- | --- | --- |
| Tahiti | 178,100 | 4,966 | 128 |
| Iles sous-le-vent | 33,100 | 1,131 | 166 |
| Moorea | 16,200 | 440 | 22 |
| Tuamotu-Gambier | 15,800 | 612 | 9 |
| Marquises | 8,600 | 455 | 21 |
| Australes | 6,800 | 733 | 36 |

### Mathematical model

We used a compartmental mathematical model to simulate vector-borne transmission [24, 25]. Both people and mosquitoes were modelled using a susceptible-exposed-infectious-removed (SEIR) framework. This model incorporated delays as a result of the intrinsic (human) and extrinsic (vector) incubation periods (Fig. 1). Since there is evidence that asymptomatic DENV-infected individuals are capable of transmitting DENV to mosquitoes [26], we assumed the same for ZIKV: all people in the model transmitted the same, regardless of whether they displayed symptoms or were reported as cases.

**Figure 1.**
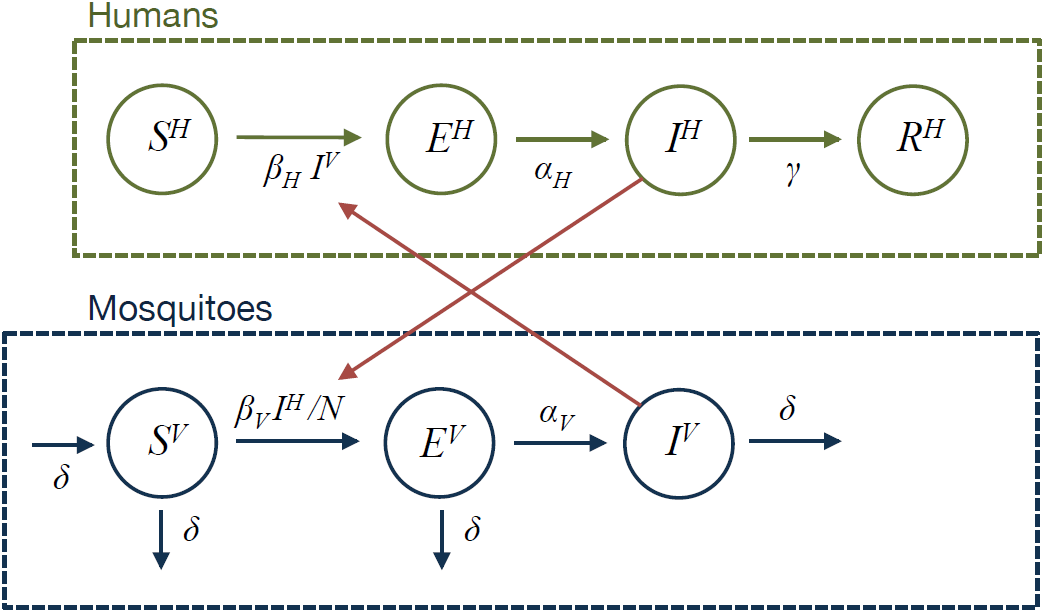
Human-vector transmission model schematic. *S*^*H*^ represents the number of susceptible people, *E*^*H*^ the number of people incubating the virus, *I*^*H*^ the number of infectious people, *R*^*H*^ the number recovered people. Similarly, *S*^*V*^ represents the proportion of mosquitoes currently susceptible, *E*^*V*^ the proportion in their incubation period, and *I*^*V*^ the proportion of mosquitoes infectious. Mosquitoes are assumed to remain infectious for life. *β*_*V*_ is the transmission rate from humans to mosquitoes; *β*_*H*_ is transmission from mosquitoes to humans; 1/*α*_*H*_ and 1/*α*_*V*_ are the mean latent periods for humans and mosquitoes respectively; 1/*γ* is the mean infectious period for humans; 1/*δ* is the mean lifespan of mosquitoes; and *N* is the human population size.

The main vectors for ZIKV in French Polynesia are thought to be Ae. *aegypti* and Ae. *polynesiensis* [12]. In the southern islands, the extrinsic incubation period for Ae. *polynesiensis* is longer during the cooler period from May to September [27], which may act to reduce transmission. Moreover, temperature can also influence mosquito mortality, and hence vector infectious period [28]. However, climate data from French Polynesia [29] indicated that the ZIKV outbreaks on the six archipelagos ended before a decline in mean temperature or rainfall occurred (Figure S1). Hence it is likely that transmission ceased as a result of depletion of susceptible humans rather than seasonal changes in vector transmission. Therefore we did not include seasonal effects in our analysis.

In the model, *S*^*H*^ represents the number of susceptible people, *E*^*H*^ is the number of people currently in their incubation period, *I*^*H*^ is the number of infectious people, *R*^*H*^ is the number of people that have recovered, *C* denotes the cumulative number of people infected (used to fit the model), and *N* is the human population size. Similarly, *S*^*V*^ represents the proportion of mosquitoes currently susceptible, *E*^*V*^ the proportion in their incubation period, and *I*^*V*^ the proportion of mosquitoes currently infectious. As the mean human lifespan is much longer than the outbreak duration, we omitted human births and deaths. The full model is as follows:

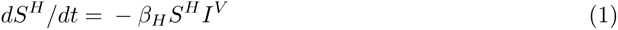

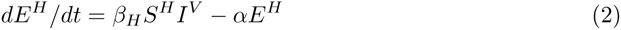

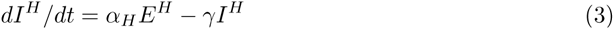

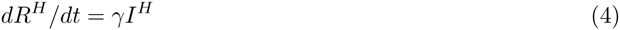

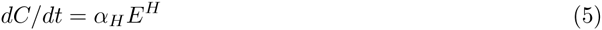

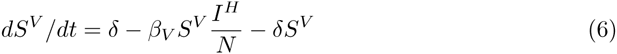

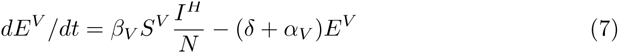

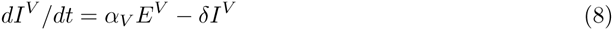

Parameter definitions and values are given in Table 2. We used weakly informative prior distributions for the human latent period, 1/*α*_*H*_, infectious period, 1/*γ*, extrinsic latent period, l/*α*_*v*_, and mosquito lifespan, 1/*μ*. For these prior distributions, we made the assumption that human latent period was equivalent to the intrinsic incubation period,i.e. that no transmission typically occurs before symptom onset. A systematic review of the incubation period for ZIKV in humans estimated a mean value of 5.9 days [30]; the infectious period, 1/*γ*, lasted for 4–7 days in clinical descriptions of 297 PCR-confirmed cases in French Polynesia [22]; the extrinsic latent period has been estimated at 10 days [1]; and mosquito lifespan in Tahiti was estimated at 10.5 days [31]. We therefore used these values for the respective means of *α*_*H*_, 1/*γ*, l/*α*_*v*_ and 1/*δ* in our prior distributions. These parameters were estimated jointly across all six regions; as mentioned above, we assumed that the parameters remained fixed over time, as temperature and rainfall levels did not change substantially during the outbreak. The rest of the parameters were estimated for each region individually; we assumed uniform prior distributions for these.

**Table 2.** Parameters used in the model. Prior distributions are given for all parameters, along with source if the prior incorporates a specific mean value. All rates are given in units of days^−1^.

| Parameter | Definition | Prior | Source |
| --- | --- | --- | --- |
| $1/\alpha_V$ | extrinsic incubation period | Gamma($\mu=10.5$, $\sigma=0.5$) | [32] |
| $1/\alpha_H$ | intrinsic incubation period | Gamma($\mu=5.9$, $\sigma=0.5$) | [30] |
| $1/\gamma$ | human infectious period | Gamma($\mu=5$, $\sigma=0.5$) | [22] |
| $1/\delta$ | mosquito lifespan | Gamma($\mu=7.8$, $\sigma=0.5$) | [31] |
| $\beta_H$ | vector-to-human transmission rate | $\mathcal{U}(0, \infty)$ | |
| $\beta_V$ | human-to-vector transmission rate | $\mathcal{U}(0, \infty)$ | |
| $r$ | proportion of cases reported | $\mathcal{U}(0, 1)$ | |
| $\phi$ | reporting dispersion | $\mathcal{U}(0, \infty)$ | |

Serological analysis of samples from blood donors between July 2011 and October 2013 suggested that only 0.8% of the population of French Polynesia were seropositive to ZIKV [33]; we therefore assumed that the population was fully susceptible initially. We also assumed that the initial number of latent and infectious people were equal (i.e.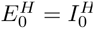), and the same for mosquitoes 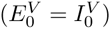. The basic reproduction number was equal to the product of the average number of mosquitoes infected by the typical infectious human, and vice versa [24]:

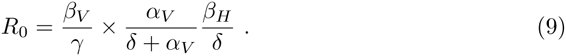

### Statistical inference

We fitted the model using Markov chain Monte Carlo (MCMC), where incidence in week *t*, denoted *c*_*t*_, was the difference in the cumulative proportion of cases over the previous week i.e. *c*_*t*_ = *C*(*t*) – *C*(*t* – 1). In the model, the total number of cases included asymptomatic and subclinical cases—which would not be detected at any site—as well as those that displayed symptoms. Hence there were two sources of potential underreporting: as a result of limited sentinel sites; and as a result of cases not seeking treatment. We adjusted for the first source of underreporting by defining *k*_*t*_ as the proportion of total sites that reported as sentinels in week *t*. We assumed that the population was uniformly distributed across the catchment areas of the healthcare sites. Under this assumption, the proportion of total sites that reported cases as sentinels in a particular week, *k*_*t*_, was equivalent to the expected fraction of new cases that would be reported in that week if the reporting proportion, *r*, was equal to 1. The parameter *r* accounted for the second source of under-reporting, and represented the proportion of cases (both symptomatic and asymptomatic) that did not seek treatment.

To calculate the likelihood of observing a particular number of cases in week *t*, *y*_*t*_, we assumed the number of confirmed and suspected cases in week *t* followed a negative binomial distribution with mean *rK*_*t*_*C*_*t*_ and dispersion parameter *ϕ*, to account for potential variability in reporting over time [34]. The dispersion parameter reflected variation in the overall proportion reported, as well as potential variation in size and catchment area of sentinel sites. Hence the log-likelihood for parameter set *θ* given data 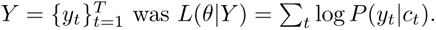 As a sensitivity analysis (see Results), we also extended the model so the likelihood included the probability of observing 314/476 seropositive individuals in Tahiti after the outbreak, given that a proportion *Z* were infected in the model. Hence for Tahiti, *L*(*θ*|*Y* = ∑_*t*_ log *P*(*y*_*t*_|*C*_*t*_) + log *P*(*X* = 314), where *X* ~ *B*(*n* = 476, *p* = *Z*). The joint posterior distribution of the parameters was obtained from eight replicates of 25,000 MCMC iterations, each with a burn-in period of 5,000 iterations (Figure S2–S7). The model was implemented in R version 3.2.3 [35] using the deSolve package [36].

### Demographic model

We implemented a simple demographic model to examine the replacement of the number of susceptible individuals over time. In 2014, French Polynesia had a birth rate of *b*=15.47 births/1,000 population, a death rate of *d*=4.93 deaths/1,000 population, and net migration rate of *m*=–0.87 migrants/1,000 [37]. The number of susceptible individuals in year *τ*, *S*(*τ*), and total population size, *N* (*τ*), was therefore expressed as the following discrete process:

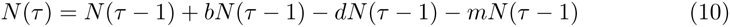

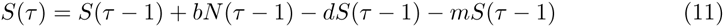

We set *S*(2014) as the fraction of the population remaining in the *S* compartment at the end of the 2013–14 ZIKV outbreak, and propagated the model forward to estimate susceptibility in future years. The effective reproduction number, *R*_eff_(*τ*), in year *τ* was the product of the estimated basic reproduction number, and the proportion of the population susceptible: *R*_eff_(*τ*) = *R*_0_*S*(*τ*). We sampled 5,000 values from the estimated joint posterior distributions of *S*(2014) and *R*_0_ to obtain the curves shown in Figure 3.

**Figure 2.**
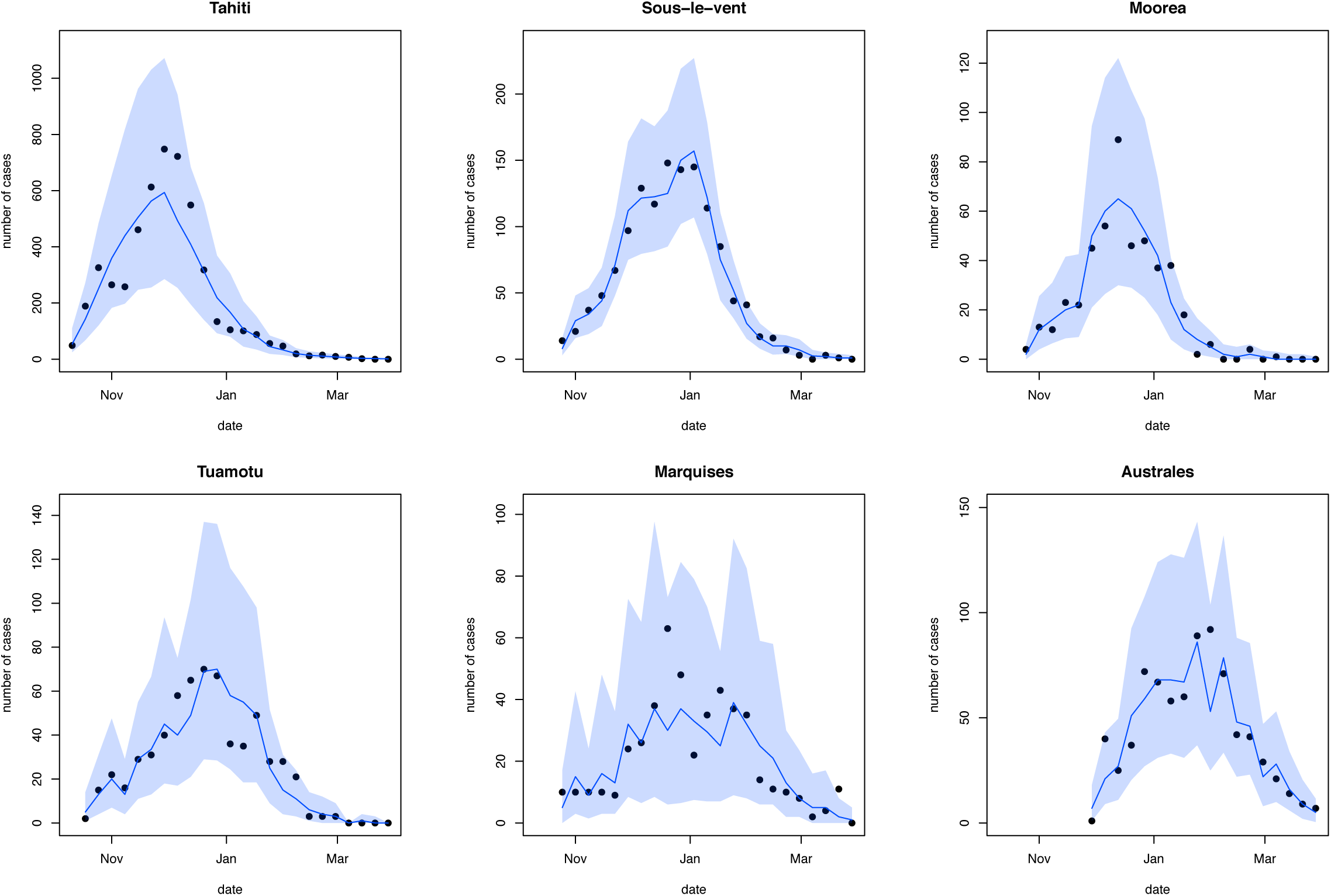
Comparison of reported cases and fitted model trajectories. Black dots show weekly reported confirmed and suspected ZIKV cases from sentinel sites. Blue line shows median of 2,000 simulated trajectories from the fitted model, adjusted for variation in reporting over time; shaded region shows 95% credible interval.

**Figure 3.**
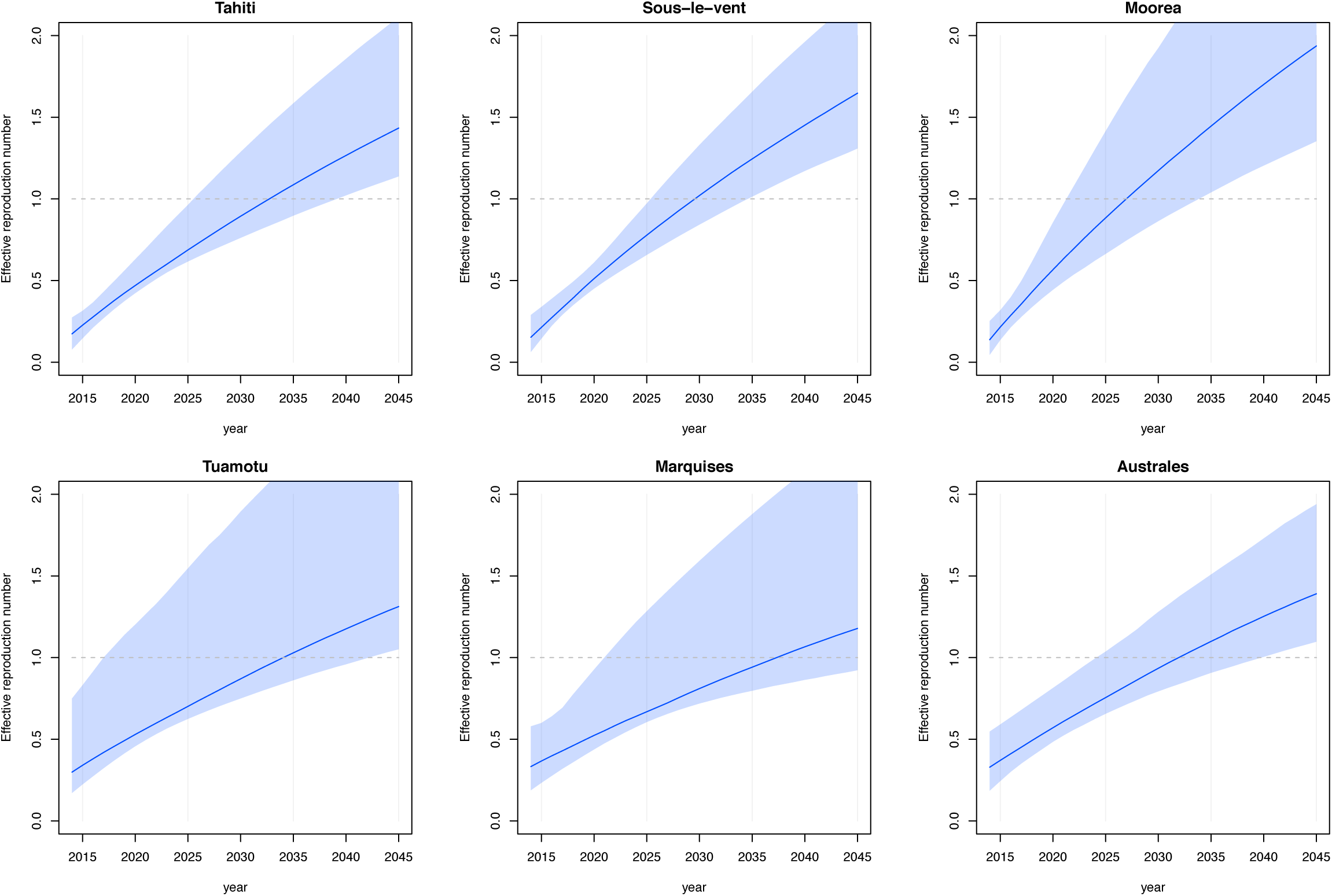
Estimated growth in effective reproduction number as susceptible pool increases over time. (A) Tahiti, (B) Sous-le-vent, (C) Moorea, (D) Tuamotu, (E) Marquises, (F) Australes. Line shows median from 1,000 samples of the posterior distribution, shaded region shows 95% credible interval.

## Results

### Epidemiological parameter estimates

Across the six regions, estimates for the basic reproduction number, *R_0_*, ranged from 2.6 (95% CI: 1.7–5.3) in Marquises to 4.8 (95% CI: 3.2–8.4) in Moorea (Table 3). Our results suggest that only a small proportion of ZIKV infections were reported as suspected cases: sampling from the fitted negative binomial reporting distributions for each region implied that 11.5% (95% CI: 7.32–17.9%) of infections were reported overall. Estimated dispersion in reporting was greatest for Marquises (Table S1), reflecting the variability in the observed data (Figure 2), even after adjusting for variation in the number of sentinel sites. Dividing the 8,744 cases reported at sentinel sites by the total estimated infections, we also estimated that 3.41% (95% CI: 3.32–3.55%) of total infections were reported at the subset of health sites that acted as sentinel sites.

**Table 3.** Estimated parameters for ZIKV infection. Estimates for the basic reproduction number, *R*_0_; the proportion of infected individuals that were reported as suspected cases at all sites (with reports following a negative binomial distribution with reporting proportion *r* and dispersion parameter *ϕ*); the total proportion of the population infected, including both symptomatic and asymptomatic cases; and the initial number of infectious humans, 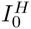. Median estimates are given, with 95% credible intervals in parentheses. The full posterior distributions are shown in Figure S2–S8.

| Region | $R_0$ | Reported (%) | Infected (%) | $I_0^H$ |
| --- | --- | --- | --- | --- |
| Tahiti | 3.5 (2.6–5.3) | 11 (5.5–20) | 95 (90–98) | 450 (71–3500) |
| Sous-le-vent | 4.1 (3.1–5.7) | 11 (7.6–15) | 96 (92–99) | 82 (3–430) |
| Moorea | 4.8 (3.2–8.4) | 7 (3.6–12) | 97 (93–99) | 58 (11–220) |
| Tuamotu-Gambier | 3 (2.2–6.1) | 6.9 (3–13) | 90 (82–96) | 92 (12–510) |
| Marquises | 2.6 (1.7–5.3) | 9.5 (2.7–23) | 87 (71–94) | 64 (13–370) |
| Australes | 3.1 (2.2–4.6) | 17 (8.2–30) | 89 (79–96) | 41 (5–140) |

### Sensitivity analyses

Our posterior estimates for the latent and infectious periods in humans and mosquitoes were consistent with the assumed prior distributions (Figure s2), suggesting either that there was no strong evidence that these parameters had a different distribution, or that the model had limited ability to identify these parameters from the available data. As a sensitivity analysis, we therefore considered two alternative prior distributions for the incubation and infectious periods for humans and mosquitoes. First, we examined a broader prior distribution. We used the same mean values for the Gamma distributions specified in Table 2, but with *σ*=2. These priors produced similar estimates for *R*_0_, proportion reported, and total number of infections (Table S2), although the estimated parameters for humans were further from zero than in the prior distribution (Figure S9).

As a second sensitivity analysis, we used prior distributions with mean values as given in studies of dengue fever, and *σ*=0.5. As there is evidence that human-to-mosquito transmission can occur up to 2 days before symptom onset [38], and the intrinsic incubation period for DENV infection is 5.9 days [39], we assumed a mean latent period of 5.9–2=3.9 days. We also assumed an infectious period of 5 days [38]; an extrinsic latent period of 15 days [39]; and a longer mosquito lifespan of 14 days [28]. Again, these assumptions produced similar estimates for key epidemiological parameters (Table S3), with posterior estimates tracking the prior distributions (Figure S10).

The estimated proportion of the population that were infected during the outbreak (including both reported and unreported cases) was above 85% for all six regions (Table 3), and we estimated that 94% (95% CI: 91–97%) of the total population were infected during the outbreak. A serological survey following the French Polynesia ZIKV outbreak found 314/476 children aged 6–16 years in Tahiti were positive for ZIKV in an indirect ELISA test for IgG antibody, corresponding to an attack rate of 66% (95% CI: 62–70) [17]. To test whether this seroprevalence data could provide additional information about the model parameters, we extended the model to calculate the likelihood of observing 314/476 seropositive individuals in Tahiti after the outbreak, as well as the observed weekly case reports. We obtained a much lower *R*_0_ estimate for Tahiti, but similar results for other regions, and the median reporting rate remained unchanged for all areas (Table S4). However, the model was unable to reproduce the Tahiti epidemic curve when the overall attack rate was constrained to be consistent with the results of the serological survey (Figure S10).

### Guillain-Barré Syndrome incidence

During the 2013–14 outbreak in French Polynesia, there were 42 reported cases of GBS [13]. This corresponds to an incidence rate of 15.3 (95% binomial CI: 11.0–20.7) cases per 100,000 population, whereas the established annual rate for GBS is 1–2 cases per 100,000 [10]. In total, there were 8,744 confirmed and suspected ZIKV cases reported at sentinel sites in French Polynesia, which gives an incidence rate of 480 (95% CI: 346–648) GBS cases per 100,000 suspected Zika cases reported at these sites. However, when we calculated the GBS incidence rate per estimated total ZIKV cases, using the model estimates based on the prior distributions in Table 2, we obtained a rate of 16.4 (95% CI: 11.5–21.4) per 100,000 cases. These credible intervals overlap substantially with the above incidence rate calculated with population size as the denominator, indicating that the two rates are not significantly different.

### Time to re-invasion

Using a demographic model we also estimated the potential for ZIKV to cause a future outbreak in French Polynesia. We combined our estimate of the proportion of the population that remained susceptible after the 2013–14 outbreak and *R*_0_ with a birth-death-migration model to estimate the effective reproduction number, *R*_eff_, of ZIKV in future years. If *R*_eff_ is greater than one, an epidemic would be possible in that location. Assuming that ZIKV infection confers lifelong immunity against infection with ZIKV, our results suggest that it would likely take 12–20 years for the susceptible pool in French Polynesia to be sufficiently replenished for another outbreak to occur (Figure 3). This is remarkably similar to the characteristic dynamics of DENV in the Pacific island countries and territories, with each of the four DENV serotypes re-emerging in sequence every 12–15 years, likely as a result of the gradual accumulation of susceptible individuals due to births [19, 40].

## Discussion

Using a mathematical model of ZIKV transmission, we analysed the dynamics of infection during the 2013–14 outbreak in French Polynesia. In particular, we estimated key epidemiological parameters, such as the basic reproduction number, *R*_0_, and the proportion of infections that were reported. Across the six regions, our median estimates suggest that between 7–17% of infections were reported as suspected cases. This does not necessarily mean that the non-reported cases were asymptomatic; individuals may have had mild symptoms and hence did not enter the healthcare system. For example, although the attack rate for suspected ZIKV disease cases was 2.5% in the 2007 Yap ZIKV outbreak, a household study following the outbreak found that around 19% of individuals who were seropositive to ZIKV had experienced ZIKV disease-like symptoms during the outbreak period [2].

Our median estimates for *R*_0_ ranged from 2.6–4.8 across the six main archipelagos of French Polynesia, and as a result the median estimates of the proportion of the populations that became infected in our model spanned 87–97%. This is more than the 66% (95% CI: 62–70%) of individuals who were found to be seropositive to ZIKV in a post-outbreak study in Tahiti. When we constrained the model to reproduce this level of seroprevalence as well as the observed weekly reports, however, we obtained a much poorer fit to the case time series (Fig. S11). The discrepancy may be the result of population structure, which we did not include within each region; we used a homogeneous mixing model, in which all individuals had equal chance of contact. In reality, there may be spatial heterogeneity in transmission, leading to a depletion of the susceptible human pool in some areas but not in others. As we used a deterministic model, differences in the estimate for reporting dispersion for different regions may to some extent reflect the limitations of the model in capturing observed transmission patterns, as well as true variability in reporting. Alternatively, it may be that not everyone who was exposed to ZIKV infected mosquitoes—which would have led to infection in the model—actually seroconverted to ZIKV after the outbreak.

The ZIKV outbreak in French Polynesia coincided with a significant increase in Guillain-Barré syndrome (GBS) incidence [13]. We found that although there was a raw incidence rate of 480 (95% CI: 346–648) GBS cases per 100,000 suspected ZIKV cases reported, the majority of the population was likely to have been infected during the outbreak, and therefore the rate per infected person was similar to the overall rate per capita. This could have implications for the design of epidemiological studies to examine the association between ZIKV infection and neurological complications in island populations.

If infection with ZIKV confers lifelong immunity, we found it would take at least a decade before re-invasion were possible. In the Pacific island countries and territories, replacement of DENV serotypes occurs every 4–5 years [19, 40], and therefore each specific serotype re-emerges in a 12–20 year cycle. The similarity of this timescale to our results suggest that ZIKV may exhibit very similar dynamics to DENV in island populations, causing infrequent, explosive outbreaks with a high proportion of the population becoming infected. In September 2014, Chikungunya virus (CHIKV) caused a large outbreak in French Polynesia [41], and is another example of a self-limiting arbovirus epidemic in island populations [5]. However, it remains unclear whether ZIKV could become established as an endemic disease in larger populations, as DENV and CHIKV have.

For immunising infectious diseases, there is typically a ‘critical community size’, below which random effects frequently lead to disease extinction, and endemic transmission cannot be sustained [18, 42]. Analysis of dengue fever outbreaks in Peru from 1994–2006 found that in populations of more than 500,000 people, dengue was reported in at least 70% of weekly records [43]. Large cities could have the potential to sustain other arboviruses too, and understanding which factors—from population to climate—influence whether ZIKV transmission can become endemic will be an important topic for future research. We did not consider seasonal variation in transmission as a result of climate factors in our analysis, because all six outbreaks ended before there was a substantial seasonal change in rainfall or temperature. Such changes could influence the extrinsic incubation period and mortality of mosquitoes, and hence disease transmission. If the outbreaks had ended as a result of seasonality, rather than depletion of susceptibles, it would reduce the estimated proportion of the population infected, and shorten the time interval before ZIKV would be expected to re-emerge.

There are some additional limitations to our analysis. As we were only fitting to a single time series for each region, we also assumed prior distributions for the incubation and infectious periods in humans and mosquitoes. Sensitivity analysis on these prior distributions suggested it was not possible to fully identify these parameters from the available data. If seroprevalence data from each region were to become available in the future, it could provide an indication of how many people were infected, which may make it possible to constrain more of the model parameters, and evaluate the role of spatial heterogeneity discussed above. Such studies may require careful interpretation, though, because antibodies may cross-react between different flaviviruses [12].

Our results suggest that ZIKV transmission in island populations may follow similar patterns to DENV, generating large, sporadic outbreaks with a high degree of under-reporting. If a substantial proportion of such populations become infected during an outbreak, it may take several years for the infection to re-emerge in the same location. A high level of infection, combined with rarity of outbreaks, could also make it more challenging to investigate a potential causal link between infection and concurrent neurological complications.

## Supporting Figures

**S1 Fig.**
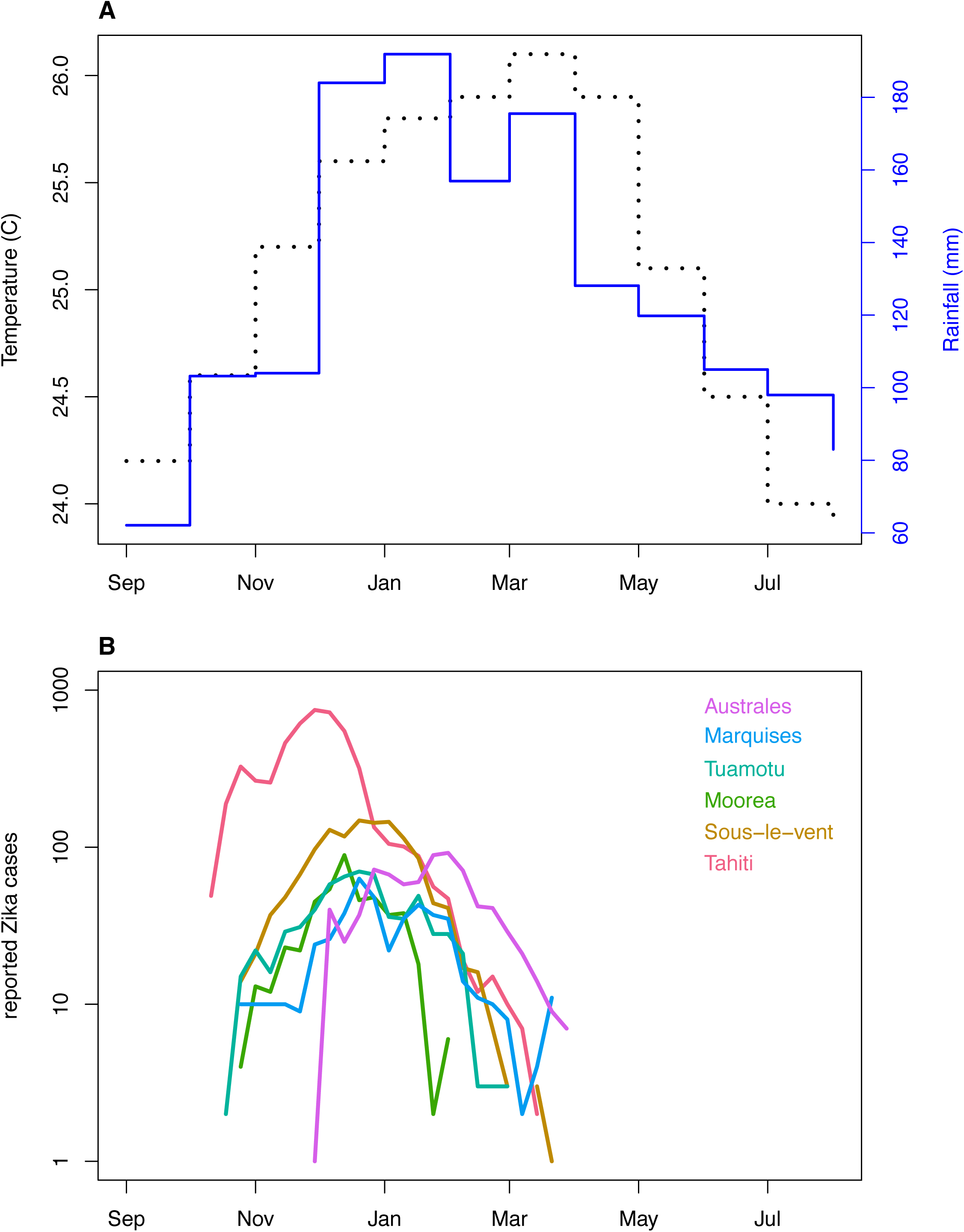
Temporal change in climate and reported Zika incidence. (A) Mean monthly temperature and rainfall in French Polynesia from 1990–2012. (B) Suspected ZIKV cases in 2013–14.

**S2 Fig.**
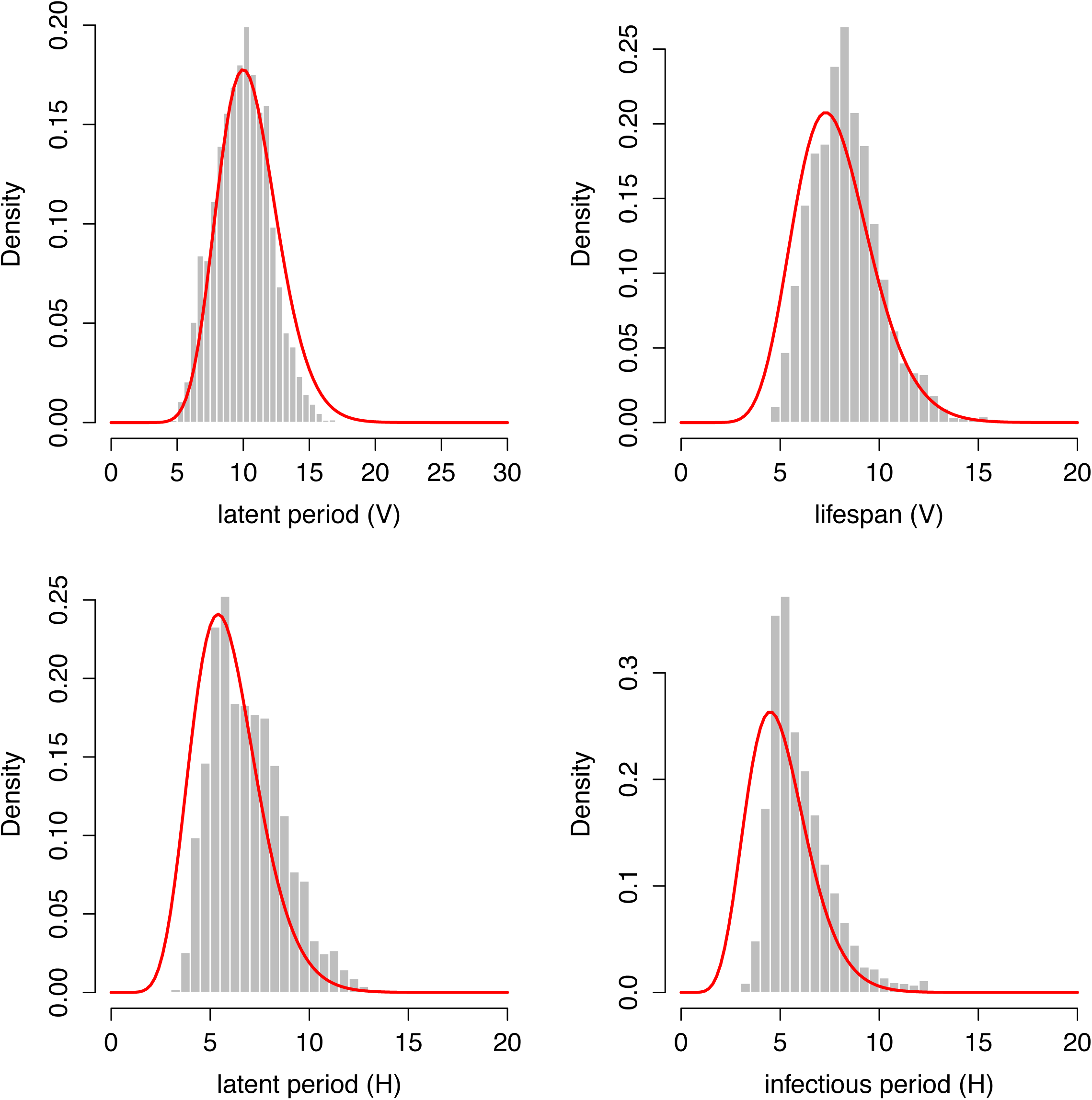
Posterior estimates for human latent period, *α*_*H*_, infectious period, 1/*γ*, extrinsic latent period, 1/*α*_*v*_, and mosquito lifespan, 1/_*μ*_. The parameters were jointly fitted across all six regions.

**S3 Fig.**
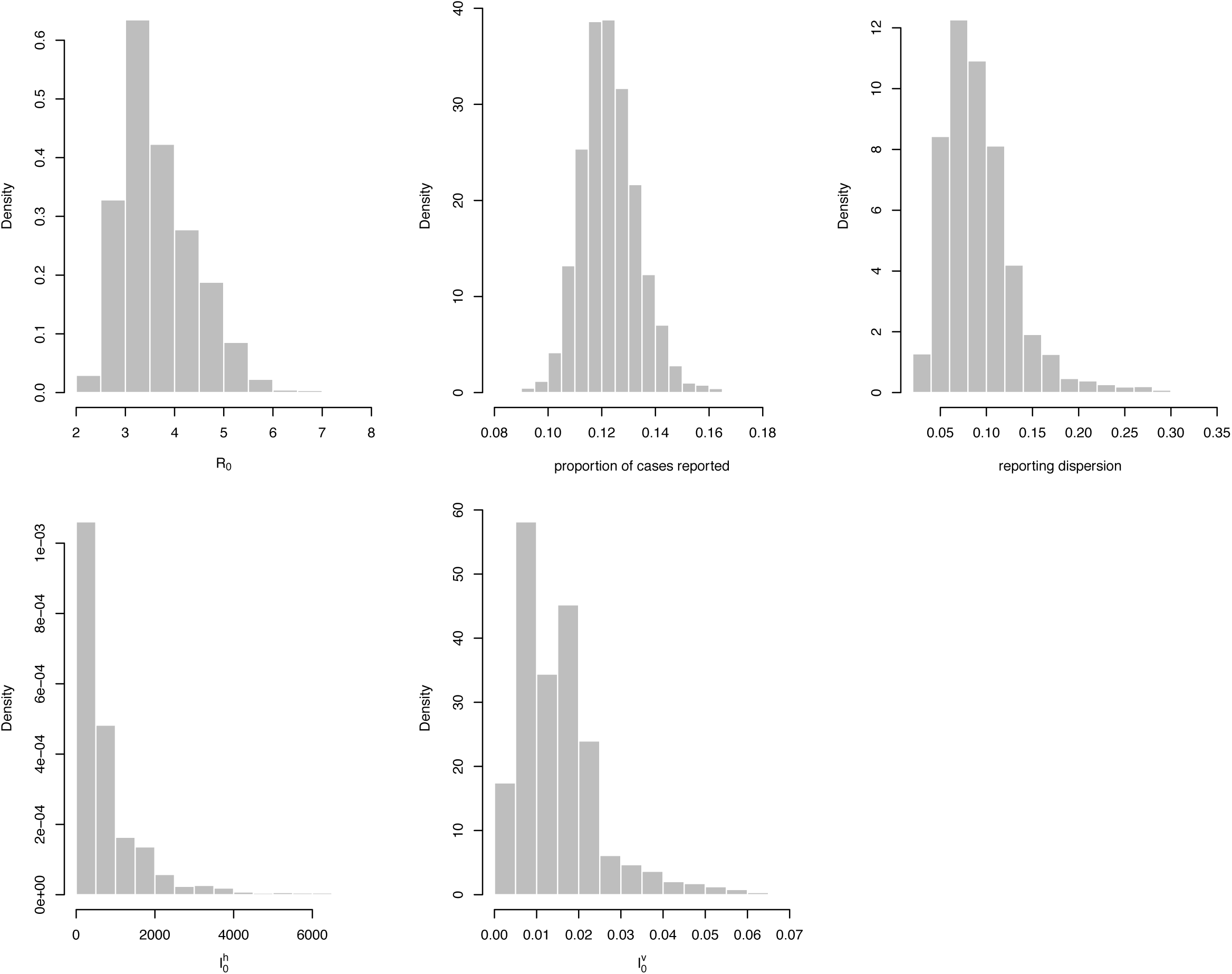
Posterior estimates for Tahiti. Plot shows marginal posterior estimates for: the basic reproduction number, *R*_0_; the proportion of cases reported, *r*; the magnitude of environmental noise, *ϕ*; the number of initially infectious humans, and the proportion of the mosquito population initially infectious.

**S4 Fig.**
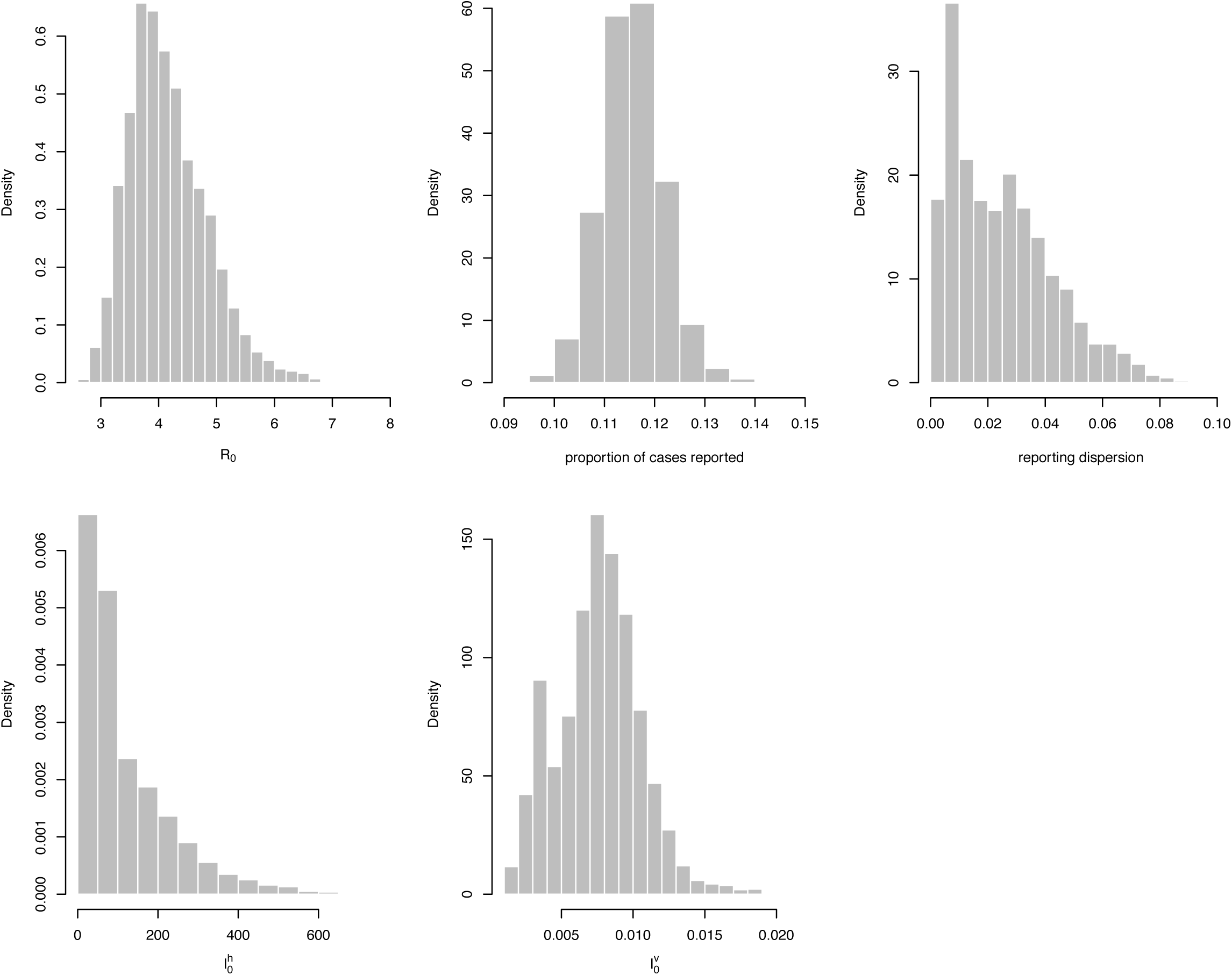
Posterior estimates for Iles sous-le-vent. Plot shows marginal posterior estimates for: the basic reproduction number, *R*_0_; the proportion of cases reported, *r*; the magnitude of environmental noise, *ϕ*; the number of initially infectious humans, and the proportion of the mosquito population initially infectious.

**S5 Fig.**
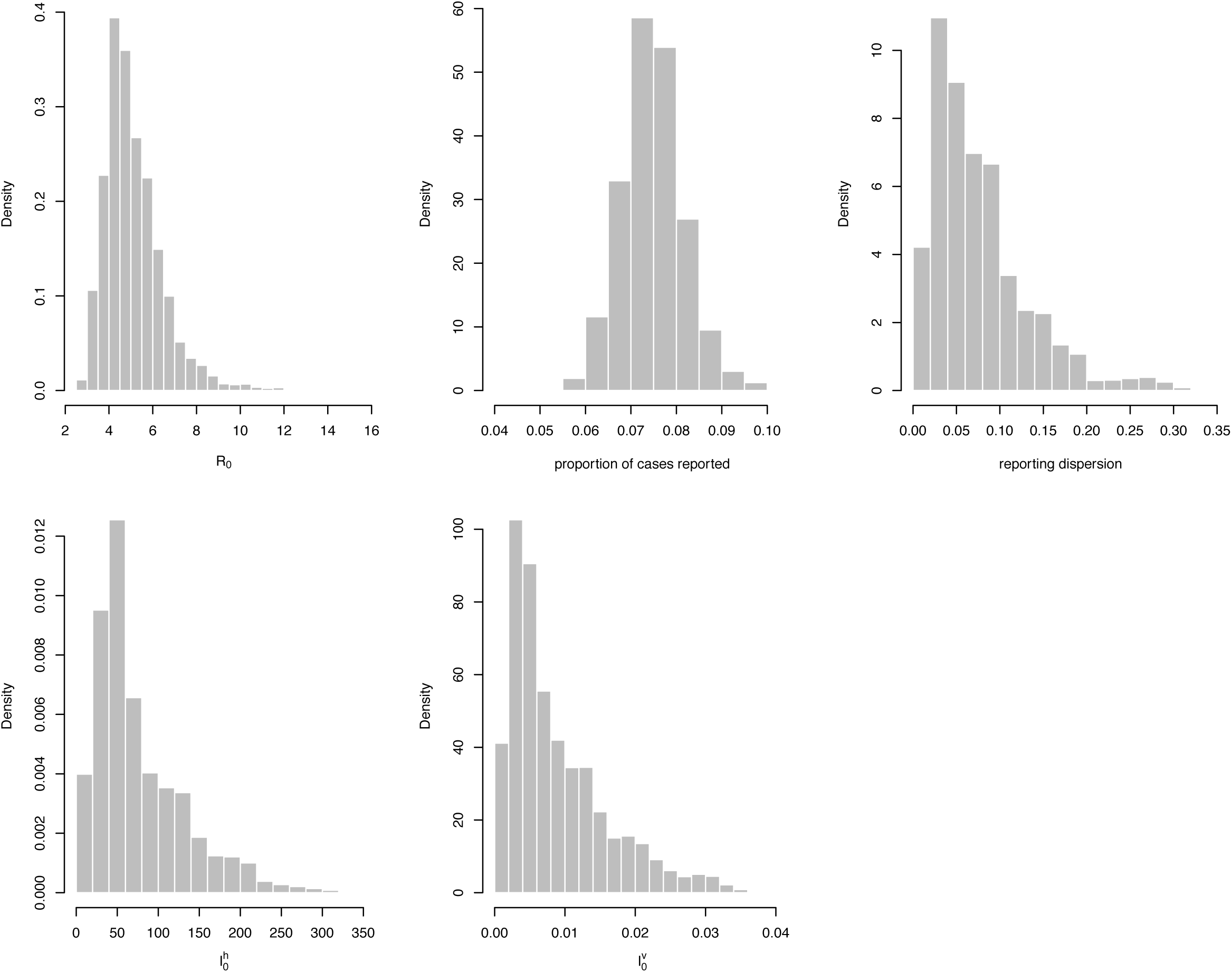
Posterior estimates for Moorea. Plot shows marginal posterior estimates for: the basic reproduction number, *R*_0_; the proportion of cases reported, *r*; the magnitude of environmental noise, *ϕ*; the number of initially infectious humans, and the proportion of the mosquito population initially infectious.

**S6 Fig.**
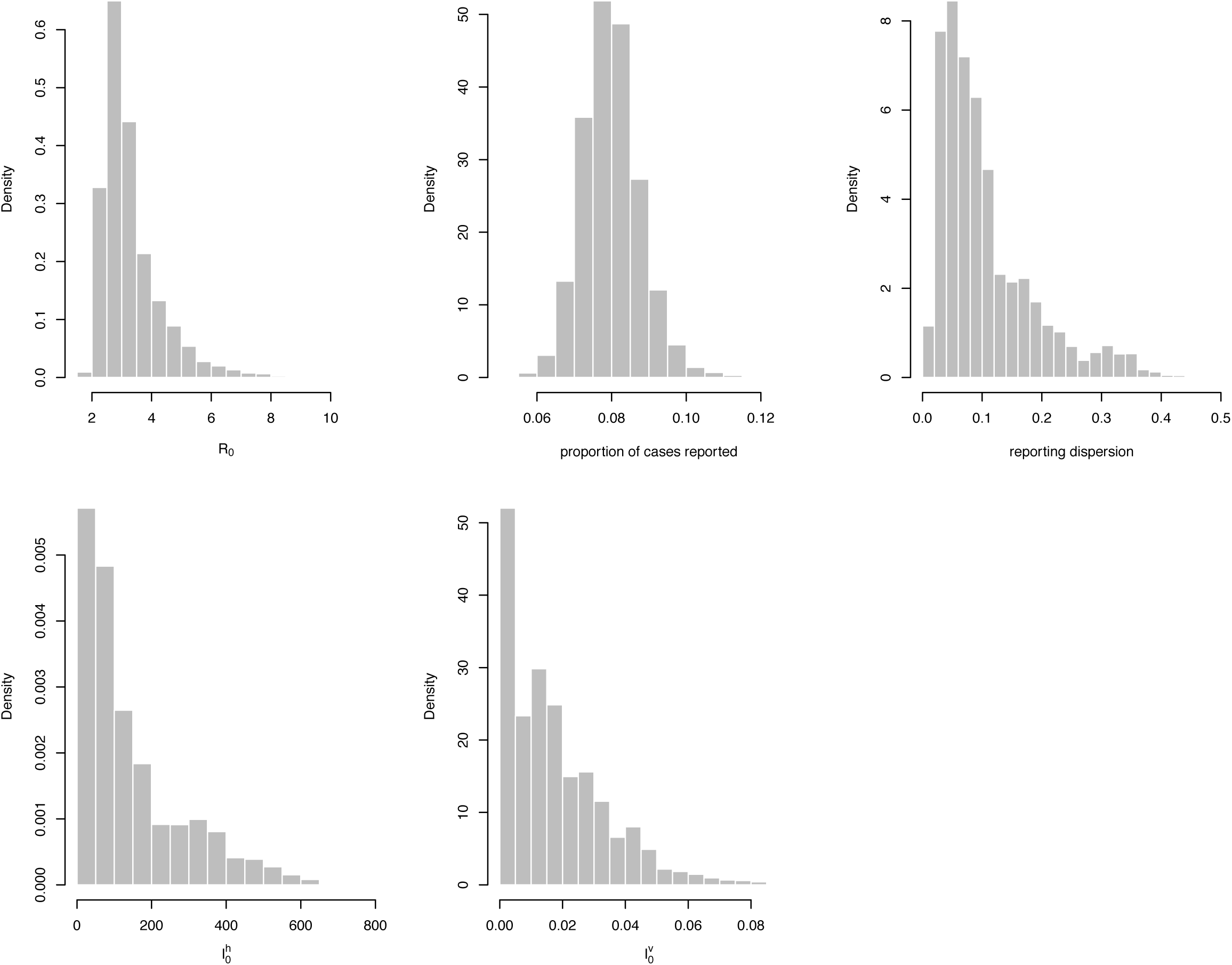
Posterior estimates for Tuamotu-Gambier. Plot shows marginal posterior estimates for: the basic reproduction number, *R*_0_; the proportion of cases reported, *r*; the magnitude of environmental noise, *ϕ*; the number of initially infectious humans, and the proportion of the mosquito population initially infectious.

**S7 Fig.**
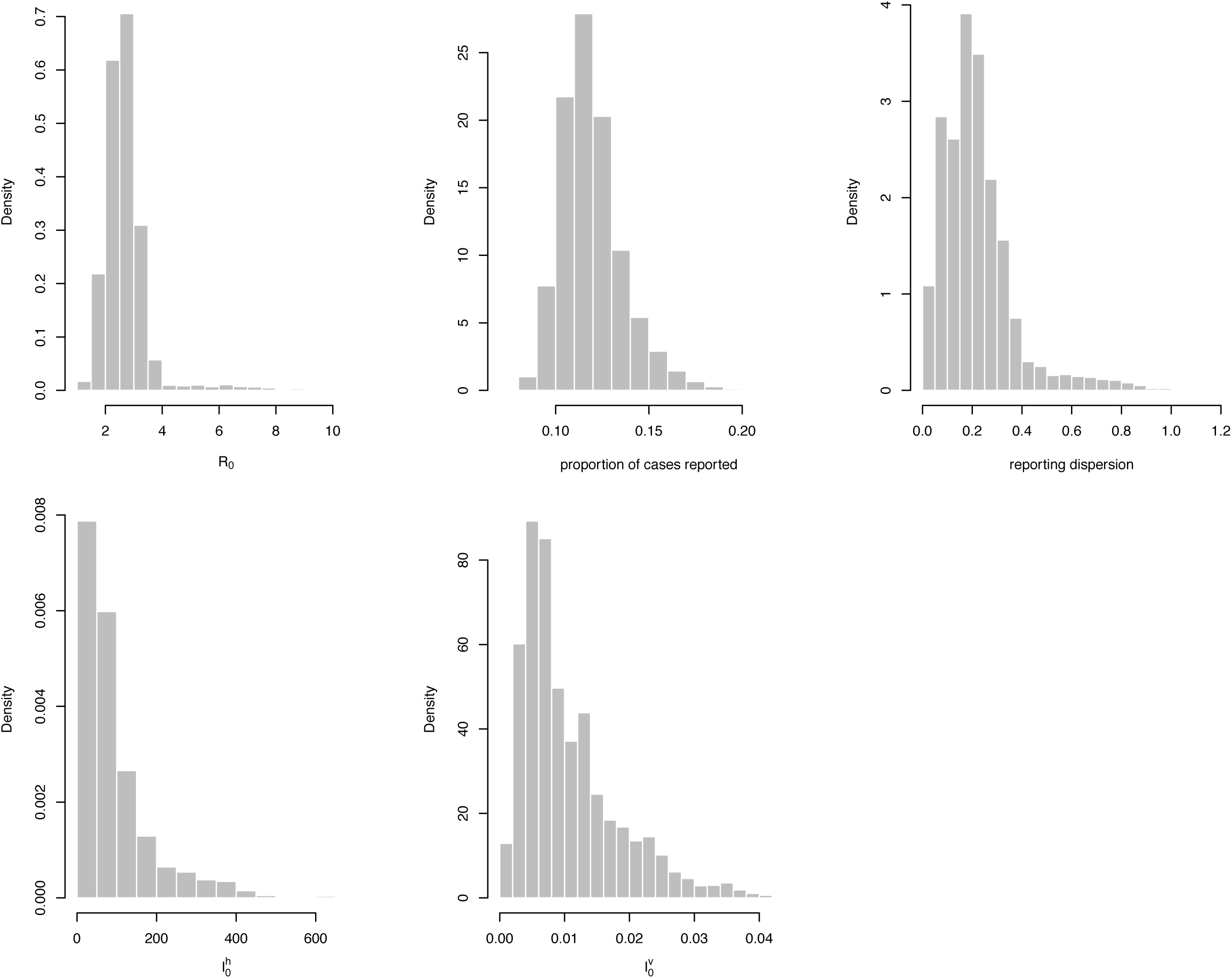
Posterior estimates for Marquises. Plot shows marginal posterior estimates for: the basic reproduction number, *R*_0_; the proportion of cases reported, *r*; the magnitude of environmental noise, *ϕ*; the number of initially infectious humans, and the proportion of the mosquito population initially infectious.

**S8 Fig.**
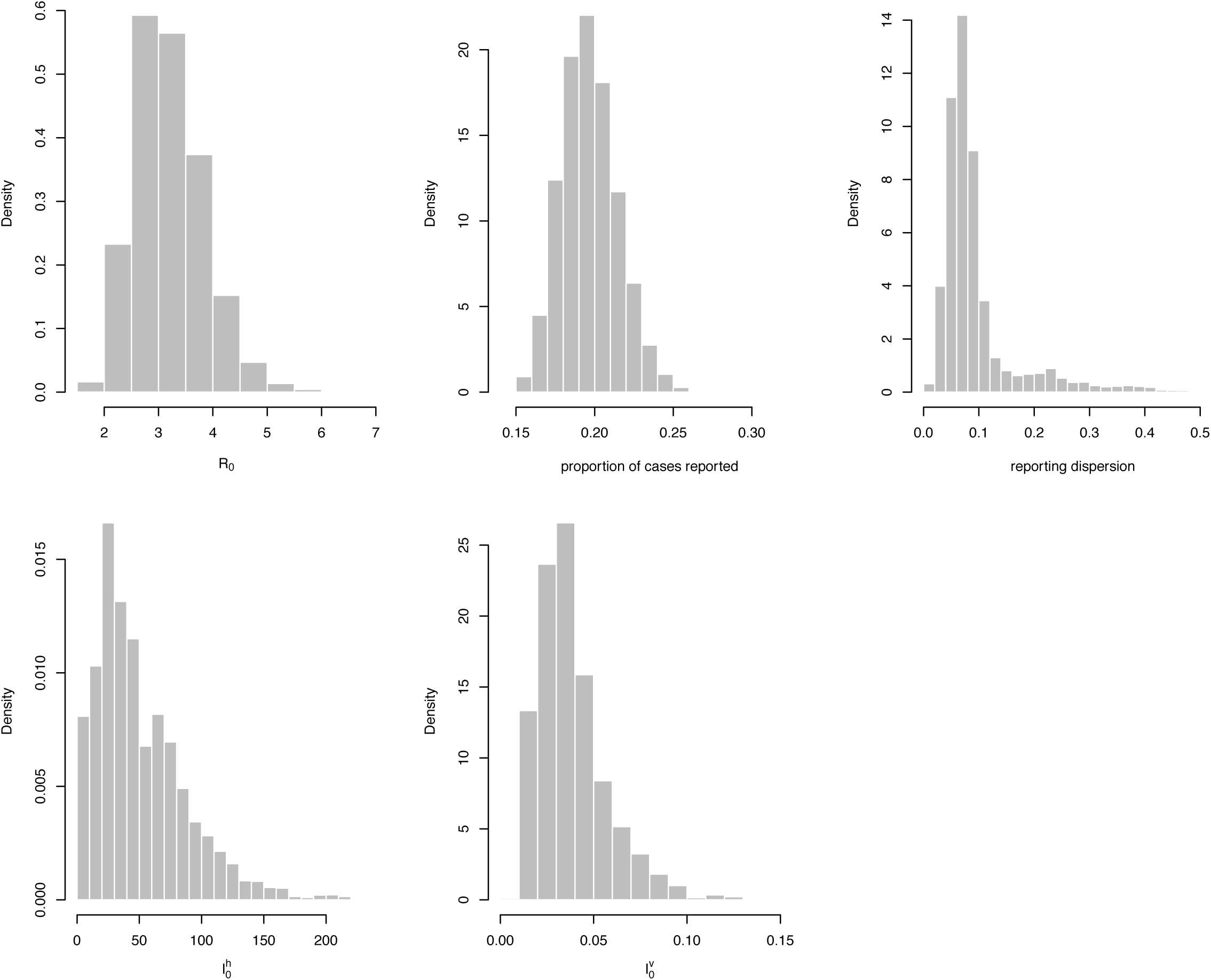
Posterior estimates for Australes. Plot shows marginal posterior estimates for: the basic reproduction number, *R*_0_; the proportion of cases reported, *r*; the magnitude of environmental noise, *ϕ*; the number of initially infectious humans, and the proportion of the mosquito population initially infectious.

**S9 Fig.**
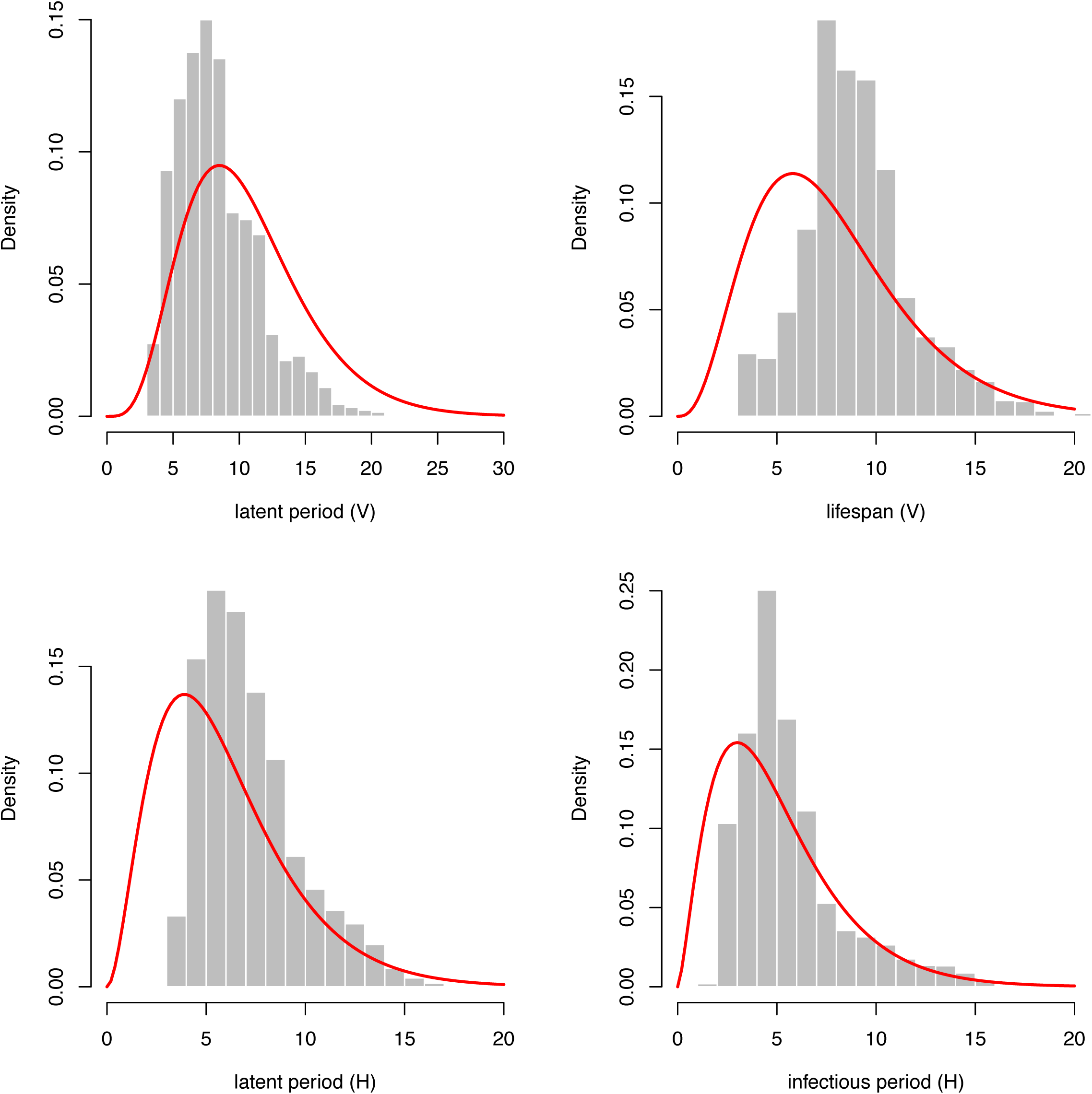
Posterior estimates when a broader prior distribution is used.

**S10 Fig.**
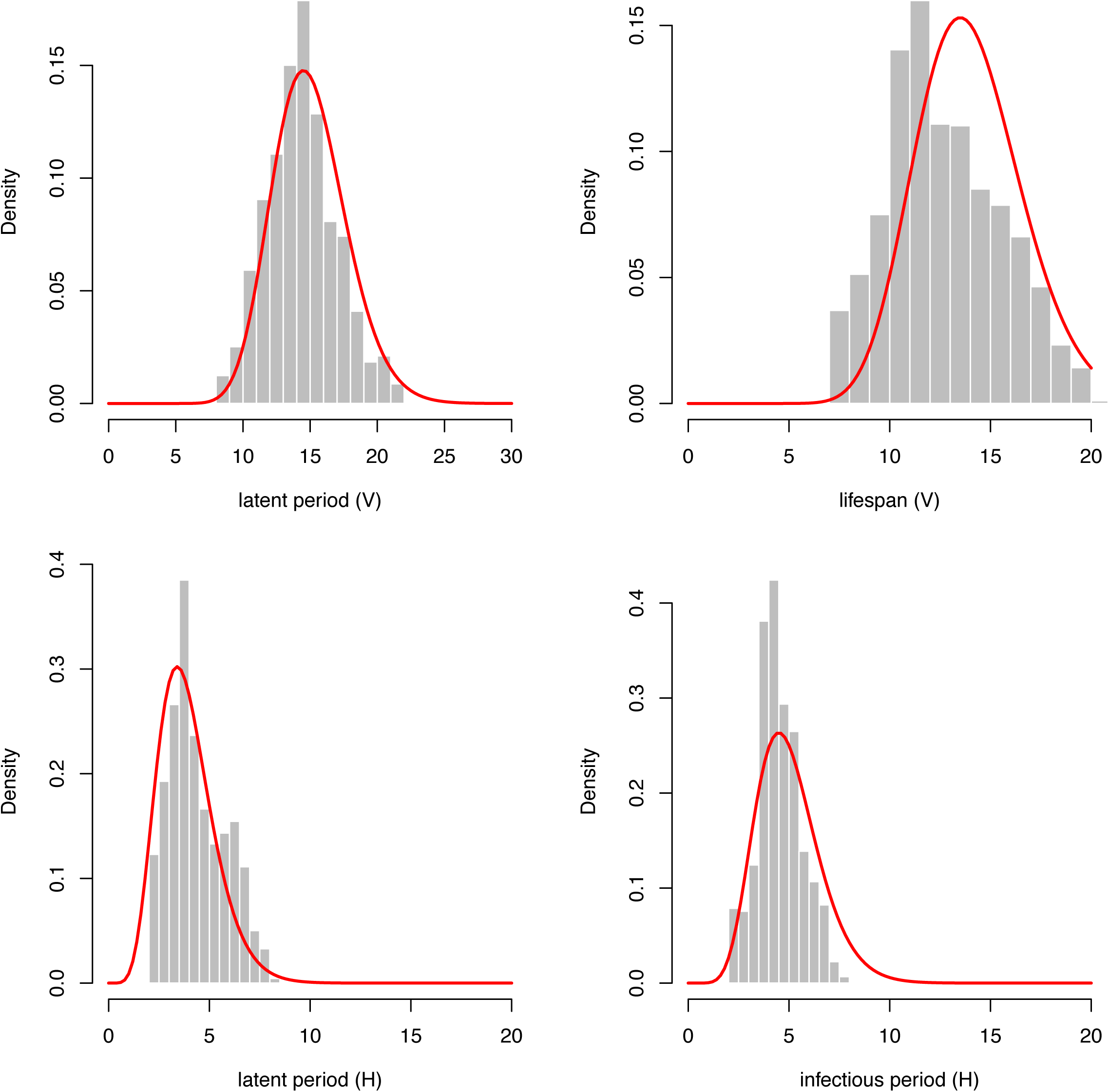
Posterior estimates when a dengue-like prior distribution is used.

**S11 Fig.**
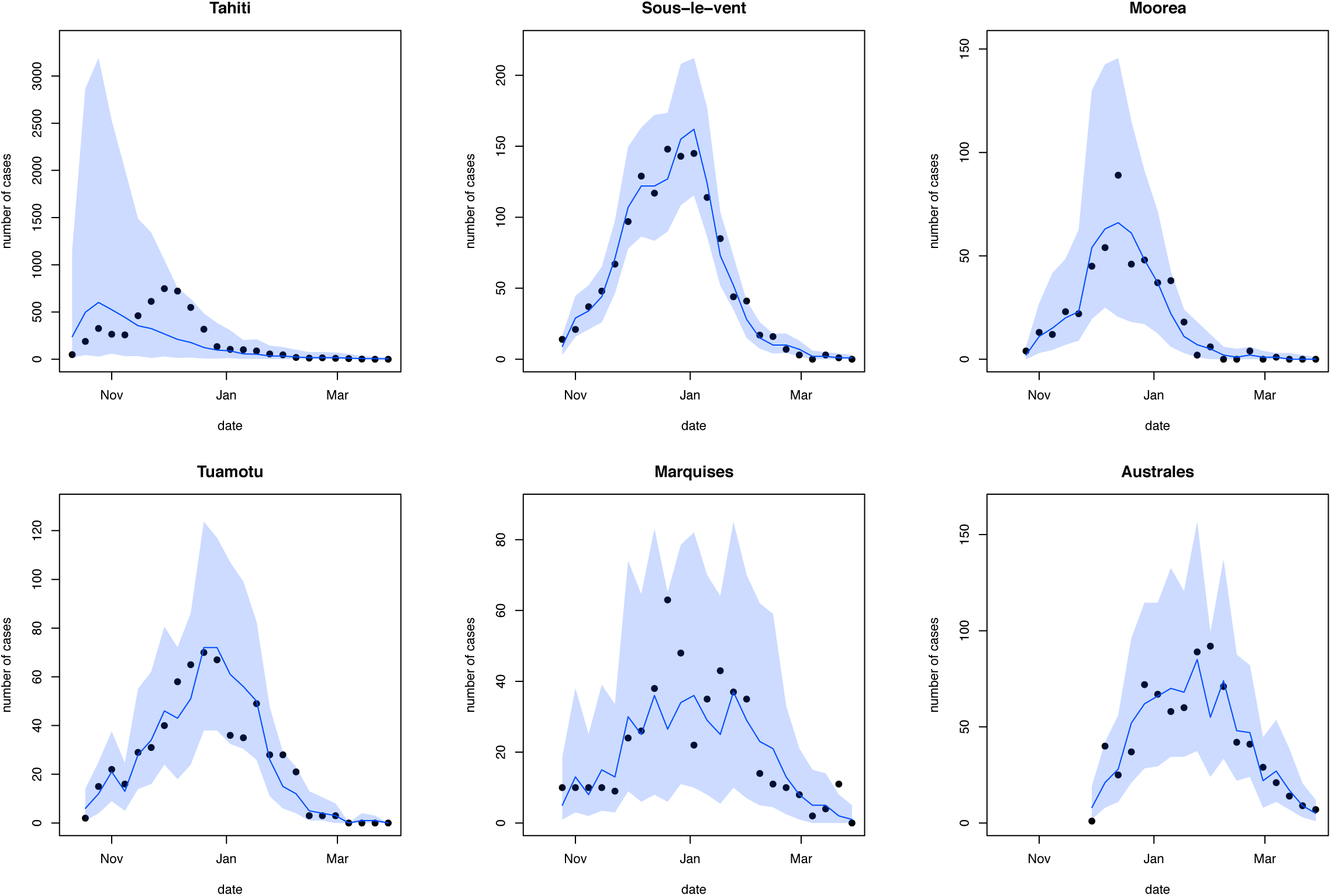
Posterior estimates when the model is fitted to seroprevalence data from Tahiti as well as the time series data.

**Table S1.**
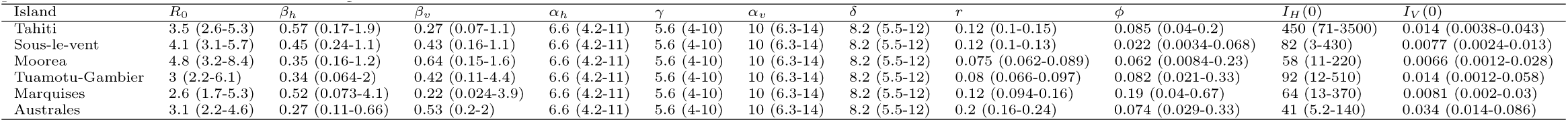
Estimated parameters for ZIKV infection. Parameters are as described in Table 2. Median estimates are given, with 95% credible intervals in parentheses. Full posterior distributions are shown in Figures S2–S8.

| Island | $R_0$ | $\beta_h$ | $\beta_v$ | $\alpha_h$ | $\gamma$ | $\alpha_v$ | $\delta$ | $r$ | $\phi$ | $I_H(0)$ | $I_V(0)$ |
| --- | --- | --- | --- | --- | --- | --- | --- | --- | --- | --- | --- |
| Tahiti | 3.5 (2.6-5.3) | 0.57 (0.17-1.9) | 0.27 (0.07-1.1) | 6.6 (4.2-11) | 5.6 (4-10) | 10 (6.3-14) | 8.2 (5.5-12) | 0.12 (0.1-0.15) | 0.085 (0.04-0.2) | 450 (71-3500) | 0.014 (0.0038-0.043) |
| Sous-le-vent | 4.1 (3.1-5.7) | 0.45 (0.24-1.1) | 0.43 (0.16-1.1) | 6.6 (4.2-11) | 5.6 (4-10) | 10 (6.3-14) | 8.2 (5.5-12) | 0.12 (0.1-0.13) | 0.022 (0.0034-0.068) | 82 (3-430) | 0.0077 (0.0024-0.013) |
| Moorea | 4.8 (3.2-8.4) | 0.35 (0.16-1.2) | 0.64 (0.15-1.6) | 6.6 (4.2-11) | 5.6 (4-10) | 10 (6.3-14) | 8.2 (5.5-12) | 0.075 (0.062-0.089) | 0.062 (0.0084-0.23) | 58 (11-220) | 0.0066 (0.0012-0.028) |
| Tuamotu-Gambier | 3 (2.2-6.1) | 0.34 (0.064-2) | 0.42 (0.11-4.4) | 6.6 (4.2-11) | 5.6 (4-10) | 10 (6.3-14) | 8.2 (5.5-12) | 0.08 (0.066-0.097) | 0.082 (0.021-0.33) | 92 (12-510) | 0.014 (0.0012-0.058) |
| Marquises | 2.6 (1.7-5.3) | 0.52 (0.073-4.1) | 0.22 (0.024-3.9) | 6.6 (4.2-11) | 5.6 (4-10) | 10 (6.3-14) | 8.2 (5.5-12) | 0.12 (0.094-0.16) | 0.19 (0.04-0.67) | 64 (13-370) | 0.0081 (0.002-0.03) |
| Australes | 3.1 (2.2-4.6) | 0.27 (0.11-0.66) | 0.53 (0.2-2) | 6.6 (4.2-11) | 5.6 (4-10) | 10 (6.3-14) | 8.2 (5.5-12) | 0.2 (0.16-0.24) | 0.074 (0.029-0.33) | 41 (5.2-140) | 0.034 (0.014-0.086) |

**Table S2.**
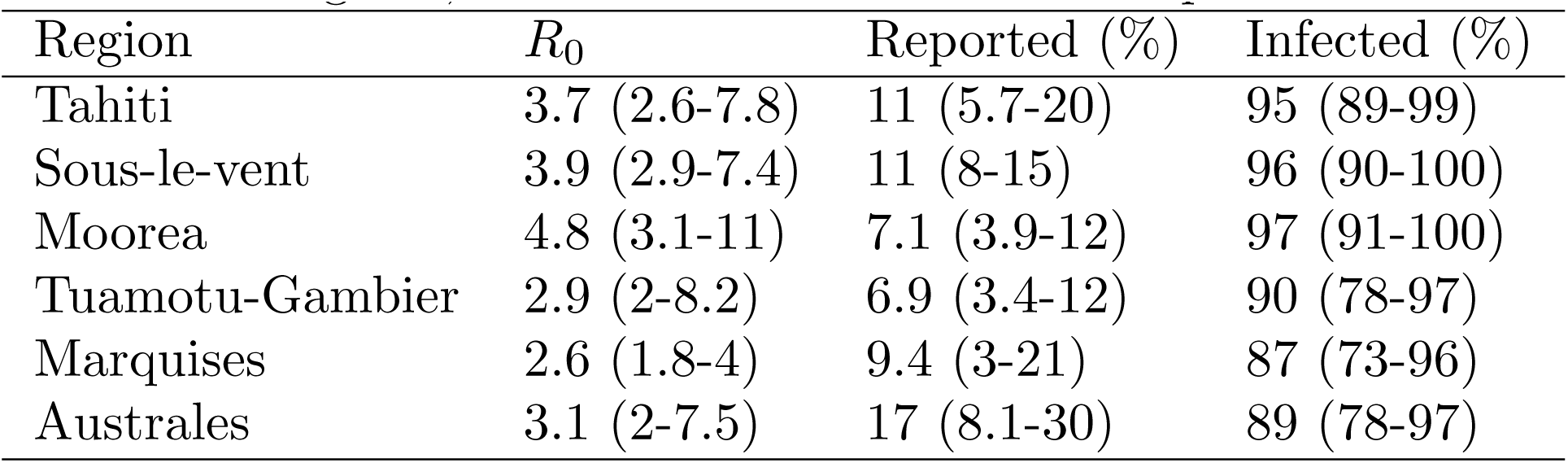
Estimated parameters for ZIKV infection when prior distributions with *σ* = 2 are used. Estimates for the basic reproduction number, *R*_0_; the proportion of infected individuals that were reported as suspected cases at sentinel sites; and the total proportion of the population infected (including both symptomatic and asymptomatic cases, with reports following a negative binomial distribution with reporting proportion r and dispersion parameter *ϕ*). Median estimates are given, with 95% credible intervals in parentheses.

| Region | $R_0$ | Reported (%) | Infected (%) |
| --- | --- | --- | --- |
| Tahiti | 3.7 (2.6-7.8) | 11 (5.7-20) | 95 (89-99) |
| Sous-le-vent | 3.9 (2.9-7.4) | 11 (8-15) | 96 (90-100) |
| Moorea | 4.8 (3.1-11) | 7.1 (3.9-12) | 97 (91-100) |
| Tuamotu-Gambier | 2.9 (2-8.2) | 6.9 (3.4-12) | 90 (78-97) |
| Marquises | 2.6 (1.8-4) | 9.4 (3-21) | 87 (73-96) |
| Australes | 3.1 (2-7.5) | 17 (8.1-30) | 89 (78-97) |

**Table S3.** Estimated parameters for ZIKV infection when dengue-like prior distributions are used. Estimates for the basic reproduction number, *R*_0_; the proportion of infected individuals that were reported as suspected cases at sentinel sites; and the total proportion of the population infected (including both symptomatic and asymptomatic cases, with reports following a negative binomial distribution with reporting proportion r and dispersion parameter *ϕ*). Median estimates are given, with 95% credible intervals in parentheses.

| Region | $R_0$ | Reported (%) | Infected (%) |
| --- | --- | --- | --- |
| Tahiti | 4 (3-6) | 11 (5.8-20) | 96 (92-98) |
| Sous-le-vent | 4.3 (3.3-5.9) | 11 (8.3-14) | 97 (92-99) |
| Moorea | 4.8 (3.4-8.3) | 7 (3.8-12) | 98 (94-99) |
| Tuamotu-Gambier | 3.4 (2.4-6.4) | 7 (3.7-12) | 91 (84-96) |
| Marquises | 2.8 (2-4.1) | 9.5 (3.3-21) | 89 (76-95) |
| Australes | 3.4 (2.4-5.3) | 17 (8.3-29) | 90 (81-95) |

**Table S4.** Estimated parameters for ZIKV infection when Tahiti serological survey is included in the likelihood. Estimates for the basic reproduction number, *R*_0_; the proportion of infected individuals that were reported as suspected cases at sentinel sites; and the total proportion of the population infected (including both symptomatic and asymptomatic cases, with reports following a negative binomial distribution with reporting proportion r and dispersion parameter *ϕ*). Median estimates are given, with 95% credible intervals in parentheses.

| Region | $R_0$ | Reported (%) | Infected (%) |
| --- | --- | --- | --- |
| Tahiti | 1.2 (0.52-1.5) | 11 (0.79-46) | 67 (63-71) |
| Sous-le-vent | 3.7 (3-5.8) | 11 (8.2-14) | 96 (92-98) |
| Moorea | 5.6 (3.1-9.9) | 7.1 (2.9-13) | 97 (92-99) |
| Tuamotu-Gambier | 2.9 (2.2-5) | 7 (4-11) | 90 (81-95) |
| Marquises | 2.6 (1.9-3.7) | 9.4 (2.8-22) | 87 (75-95) |
| Australes | 3.1 (2-4.9) | 17 (8.7-30) | 89 (77-95) |

